# The critical role of mental imagery in human emotion: insights from Aphantasia

**DOI:** 10.1101/726844

**Authors:** Marcus Wicken, Rebecca Keogh, Joel Pearson

## Abstract

One proposed function of imagery is to make thoughts more emotionally evocative through sensory simulations. Here we report a novel test of this theory utilizing a special population with no visual imagery: Aphantasia. After using multi-method verification of aphantasia, we show that this condition, but not the general population, is associated with a flat-line physiological response to frightening written, but not perceptual scenarios, supporting imagery’s critical role in emotion.

---

Models of human cognition propose mental imagery exists for aiding thought predictions by linking them to emotions^1 2^. For example, models of mental illness commonly identify mental imagery as a key driver of negative emotion ^2 3 4^, the sensory simulation of non-current events is hypothesised to drive affective reactions to such thoughts.

However, theories proposing imagery exists to link thoughts with an emotional response are largely based on subjective reports of emotions from individuals attempting to think with and without sensory simulations.^5^ It is likely that for most of us, sensory simulations in thought are largely involuntarily^6^, and hence cannot be switched-off on cue.

Here, we tested the theory that imagery can function to enhance emotions in thought, by utilizing a group of individuals who do not experience visual imagery, a naturally occurring ‘knock-out’ model of image*less* cognition. The subjective absence of visual mental imagery ability in such individuals has been documented since the 19^th^ century,^7^ however the condition was only recently given a name, aphantasia^8^, and has to date been defined by subjective reports and floor scores on the self-report Vividness of Visual Imagery Questionnaire (VVIQ^9^).

Self-identified aphantasics approached the lab, wherein we first verified their lack of imagery using the VVIQ (see **Supplementary Figure S1**; score cut-off ≤32^10^). However, were we to solely rely on self-report measures such as the VVIQ, self-identified aphantasics may simply lack good metacognition of their imagery or have a bias to report it as low or non-existent. We therefore further verified the absence of imagery in this population using a psychophysical task that objectively measures imagery’s sensory strength via imagery’s ability to systematically prime perception in subsequent binocular rivalry (score cut-off ≤0.60, see description of paradigm in **Supplementary Materials**)^11,12^. Using these criteria, a sample of 22 out of 29 aphantasics (9 female) qualified to participate in our study and their performance was compared to 24 control participants (11 female), who self-identified as having intact imagery and had VVIQ scores in excess of the cut-off. Although our aphantasic group were on average older than controls (Aphantasic: mean age 33.7, SD 12.1 Control: mean age 23.0, SD 9.5) the possibility that age was a significant factor in our results was subsequently ruled out (See **Supplementary Figure 3**).

Next, to test how image-based thoughts might amplify emotions, participants read a series of custom, speed-controlled, first-person fictitious fearful scenarios (**Figure 1A**; Methods), while their skin conductance level (SCL) was continuously recorded. Skin conductance indexes changes in autonomic nervous system arousal, and generally increases in response to frightening stimuli,^13^ including imagined stimuli.^14^ Participants read the scenarios presented as a series of visually descriptive phrases on screen in a darkened room, totalling 100 seconds per story. Each participant underwent five distinct story trials (see **Methods**). Participants’ comprehension of each story was verified afterwards (see **Methods**). For each trial, skin conductance level was baseline-corrected by subtracting the mean SCL recorded during that trial’s preceding baseline period. Average baseline corrected data was then extracted from each trial in consecutive five-second time bins. **Figure 1B** shows both the aphantasic (blue) and control participant SCL data (red), during the imagery provoking scenarios. The rightmost bar plot shows the mean SCL collapsed over the full 100 seconds of the stories’ duration. A two-tailed t-test on the difference between this collapsed mean revealed a significantly higher SCL in controls relative to aphantasics, t(44) = 2.18, *p* = 0.035), suggesting that aphantasic individuals show a reduced fear response to the frightening scenarios when compared to controls. Underscoring this effect, further two-tailed t-tests revealed that only the control group experienced a significant increase in SCL compared to zero (*p* = .0004), while the aphantasic group’s increase did not significantly differ from zero (*p*=.54).

**Figure 1A.**
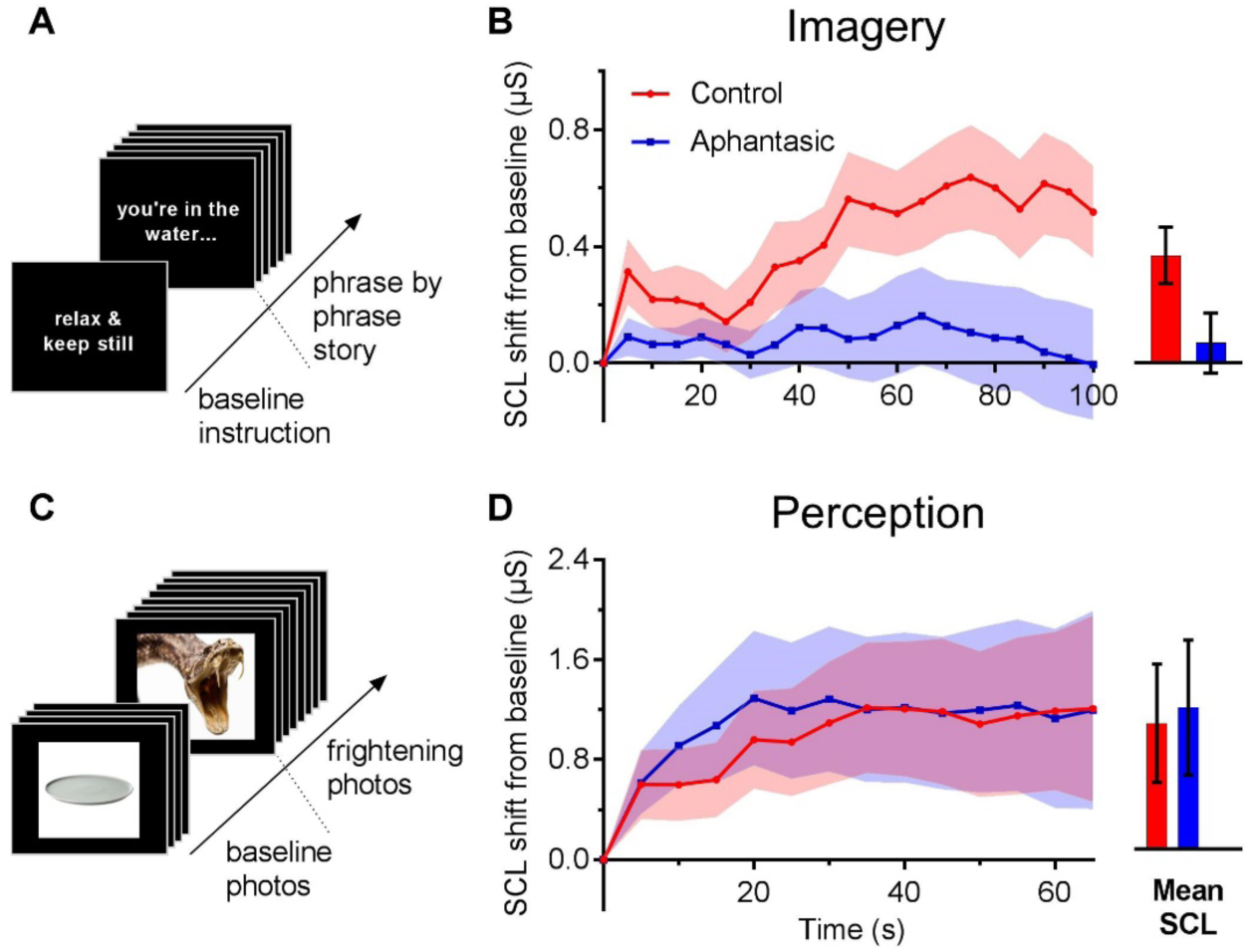
Imagery Experiment. 50s of baseline SCL was recorded prior to each scenario trial while participants viewed an on-screen instruction. Next, each scenario trial was presented to participants as a succession of 50 on-screen phrases, each displayed for 2s. **B. *Left:*** Aggregated progressions of baseline-corrected SCL across the duration of scenarios (sampled as average across 5s time bins). ***Right***: Mean and SEM across time bins. **C. Perception Experiment.** Baseline SCL was recorded while participants viewed neutral photos, before being presented with a succession of frightening photos. Photos appeared on screen for 5s each and immediately followed one another. **D. *Left:*** Aggregated progressions of baseline-corrected SCL across the duration of the frightening photos sequence (sampled as average across 5s time bins). ***Right:*** Mean and SEM across time bins.

These results suggest that the arousal response to reading the fictitious scenarios may be largely contingent on having intact imagery to simulate the scenario content. This is consistent with existing evidence for imagery’s theorized role as an emotional amplifier^15^. However, it is also possible that the difference in SCL change reflects group differences unrelated to imagery. One possibility is that our control group was more prone to general anxiety than our aphantasic group, or that aphantasic individuals were for some reason generally less emotionally responsive. However, a comparison of trait anxiety scores from the State-Trait Anxiety Inventory (STAI)^16^ suggested equivalent levels of trait anxiety between the two groups (see **Supplementary Figure S2**).

Another possibility is that aphantasic individuals have a reduced/dampened fear response in general. To investigate this non-imagery explanation of the data, we ran a follow up perception experiment (**Figures 1C and 1D**). In this experiment SCL was measured in the same manner, however participants viewed frightening perceptual stimuli, instead of reading frightening fictitious scenarios. In the Perception experiment aphantasic (n=15, 9 female), and control (n=14, 7 female) participants passively viewed a continuous presentation of 18 International Affective Picture System (IAPS)^17^ photographs (5s each).

As shown in **Figure 1C** the first five photographs served as a neutral baseline (rated least frightening in IAPS norms (e.g. umbrella, dinner plate)), while the remaining 13 were rated very frightening (e.g. snake’s mouth, assault, cadaver). The resulting data was averaged across participants within each group to again produce two plots (**Figure 1D)**. Both groups showed a monotonic increase in baseline-corrected SCL when viewing frightening perceptual pictures. Again, the rightmost bar plot shows the mean SCL collapsed over all time bins. Two-tailed t-tests revealed that both groups experienced significant increases in SCL compared to zero, while viewing the frightening pictures (aphantasic *p* = .046, control *p*= .036), and the overall magnitude of the increase did not significantly differ between aphantasics and controls, t(28) = 0.141, *p* = 0.89. These results suggest that it is not the case that aphantasic individuals have a general reduction in fear response, indicating that the observed differences in SCL is likely due to group differences in visual imagery ability.

Together, the results show that aphantasia/imagery has a marked impact on physiology when reading frightening fictitious scenarios. These data suggest that imagery connects thoughts to emotions, vis sensory simulation. This is, to the best of our knowledge, the strongest evidence to date supporting imagery’s theoretical role as an ‘emotional amplifier’. This adds further impetus to the growing clinical appreciation that the imagistic content of cognition is a central determinant of mental wellbeing, and that image-based cognition is more emotionally consequential than its propositional (e.g. verbal) counterparts, despite these often being the explicit focus of ‘talk therapy’.

These data also provide further support for aphantasia being a condition that is characterised by a true lack of visual imagery, rather than a condition of poor metacognition. The results also raise questions about the emotional consequences of being, or *not* being, aphantasic. It is important to note the results do not suggest that aphantasia is associated with general muted emotionality.

## Methods

### Skin Conductance Experiments

#### Participants

Twenty-two (nine female) self-described aphantasic participants completed all experiments and questionnaires. The Aphantasics were recruited through a Facebook page, had emailed the lab regarding their aphantasia, or were referred to the lab by Professor Adam Zeman, a UK-based aphantasia researcher. Aphantasic participants were financially reimbursed for their time. The 24 (11 female) control participants were psychology students recruited via the UNSW research participation for course credit system. All experiments were approved by the UNSW psychology ethics committee.

#### Apparatus

For both skin conductance experiments, participants were seated in a blackened, sound-attenuated room with lights off, in front of a BENQ ZL2420-B LCD computer monitor with a viewing distance of approximately 60cm. All stimuli were displayed against a black background using Matlab v.R2014b with the Psychophysics Toolbox (v.3.0.12) extension^18^ on an HP Z240 Tower Workstation computer. A second HP Z240 Tower Workstation, interfaced with the display computer, continuously recorded skin conductance level using ADInstruments hardware (Powerlab Model 16/30, MLT116F finger electrodes, and ML116 GSR Amplifier) and associated Labchart Pro v.8.1.4 software^19^ at a 1000 Hz sampling rate. The finger electrodes were attached to the distal phalanges of the participant’s non-dominant hand. The in-room air conditioning setting was maintained at 20 degrees Celsius.

#### Stimuli

All on-screen text stimuli for both experiments were displayed in white 14 point Courier New font. A text instruction reading “relax, clear your mind, and keep still” appeared during baseline skin conductance recording prior to each fictitious scenario in the Imagery experiment, and prior to presenting the photographs in the Perception experiment.

#### Procedure

##### SCL imagery task

In the Imagery experiment, fictitious scenarios were presented as a succession of 3-7 words phrases, immediately following one another, each displayed for 2 seconds. One of the five scenario scripts (Shark Attack) is provided below as an example (Forward-slashes separate on-screen phrases here and were not displayed in the experiment):

> you are at the beach / in the water / looking out to sea / blue sky, grey ocean / your hands wade / through the water / you bob in the swell / your eyes dip below / dark blue-green void / you look towards shore / people on the beach / one of them is pointing / another shields their eyes / staring towards you / you turn in the water / look back out to sea / sun glints off the waves / suddenly a dark flash / in the distant waves / maybe it was a shadow / you turn to the beach / more people are pointing / they look anxious / looking back out to sea / a large fin / slices the surface / moving closer / you back-kick towards shore / your feet foam the water / obscuring your view / you turn onto your belly / you swim for shore / you see the distant crowd / between your splashing arms / your face dips below / at right, a looking shape / approaching fast / then shooting off behind / you resurface / the fin is closer / slicing the water towards you / below it a dark blur / wide as a car / you look under water / a wide open mouth / your legs flail / before white teeth / they disappear inside / cloudy red engulfs your vision / the surface above spins out of sight.

Participants’ proper reading and comprehension of each scenario was verified by asking them to briefly summarize the events described in each scenario at the conclusion of the experiment. Participants had no prior warning that they would be asked to do this. Where a participant could not recall the events of each scenario, it was assumed that they had not adequately attended to the experiment and their data was excluded from analysis.

In the Perception experiment, IAPS^20^ images were presented in the centre of the screen in their original JPEG format and size. IAPS image catalog numbers used, and the order of presentation for all participants was as below:

> *Neutral photographs:* 7006.JPG, 7010.JPG, 7150.JPG, 7045.JPG, 7150.JPG;
>
> *Frightening photographs:* 3001.JPG, 3061.JPG, 6555.JPG, 3016.JPG, 1932.JPG, 1120.JPG, 1304.JPG, 2120.JPG, 3530.JPG, 5972.JPG, 6370.JPG, 9620.JPG, 9908.JPG.

#### Binocular rivalry sensory strength of imagery assay

Binocular rivalry occurs when two distinct stimuli are presented simultaneously, one to each eye, resulting in only one of the stimuli being perceived, while the other is suppressed from awareness. If stimuli properties are corrected to account for inconsistencies in participants’ eye dominance, then each stimulus is equally likely to be perceived during a brief rivalry display. Over viewing time, the initially perceived and suppressed stimuli tend to switch, such that perception fluctuates randomly between the two possibilities, rather than comprising a mixture of both. It is known that exposing a person to a weak perceptual display of one of the stimuli alone prior to a brief rivalry display will tend to bias their initial perception toward that weak stimulus^21^,^22^. In other words, participants will perceive the weakly displayed stimulus in the subsequent rivalry display on a higher proportion of trials than would be expected by chance (50%). A similar pattern emerges when participants imagine one of the stimuli prior to a rivalry display, leading to the conclusion that imagery acts like weak visual perception^23^. The proportion of trials in which a participant’s prior imagery matches their perception in rivalry has been used as an index of the sensory strength of that individuals’ visual mental imagery. The measure offers an objective measurement of the sensory strength of an individual’s mental imagery via its facilitative impact on visual perception, thereby bypasses the metacognitive subjectivity inherent in traditional self-report measures of imagery’s vividness, such as the VVIQ. Self-identified aphantasics underwent a version of this binocular rivalry sensory strength of imagery assay (as well as the VVIQ) prior to participating in this study. The apparatus, stimuli, and procedure are described in full in previous work^24^, and the results for participants in this study are summarized in Supplementary Figure S1.

#### Apparatus

The binocular rivalry task was completed in a completely blackened room. Participants viewing distance was fixed at 57cm using a chin rest. Participants wore red-green anaglyph glasses throughout the experiment, and the experiment was conducted on an 27 inch iMac with a resolution of 2560 x 1440 pixels, with a frame rate of 60 Hz.

#### Stimuli

The binocular rivalry stimuli consisted of red horizontal (CIE x = .57, y = .36) and green vertical (CIE x = .28, y = .63) Gabor patterns, 1 cycle/, Gaussian σ 1.5°. The patterns were presented in an annulus around the fixation point and both Gabor patterns had a mean luminance of 4.41 cdm2.

Mock rivalry images were presented on 12.5% of trials and these consisted of a physically half-half red-horizontal and green-vertical Gabor patch, consisting of the same above parameters. The mock image was split horizontally with a division that varied from trial to trial using a zig-zag random walk.

#### Binocular rivalry imagery task procedure

Participants first underwent the previously documented eye-dominance task to control for any differences in eye-dominance.^25^ Following this they completed the binocular rivalry imagery task (see S4 for experimental timeline). At the beginning of each trial participants were cued to imagine one of two images with the letter “R” (red-horizontal Gabor patch) and “G” (green-vertical Gabor patch). This was followed by a six second imagery period where the participants were asked to imagine the image in as much detail as they could. Following this a very brief (750ms) binocular rivalry display was presented and participants indicated which of the two images was dominant in the display 1 = green 2 = equally mixed 3 = red. Participants completed two blocks of 48 trials, resulting in 96 trials in total.

## Supplemental results

A potential confounding influence on differences in skin conductance changes was participant age, as recruiting limitations obstructed age-matching and aphantasics were on average older than controls. However, analysis of the relationship between skin conductance level and age within each group in the Imagery experiment revealed mild positive correlations in both cases (**Figure S3**), the opposite of what would be expected if the aphantasic group’s seniority was driving their weaker SCL response.

## Supplementary Figures

**Supplementary Figure S1.**
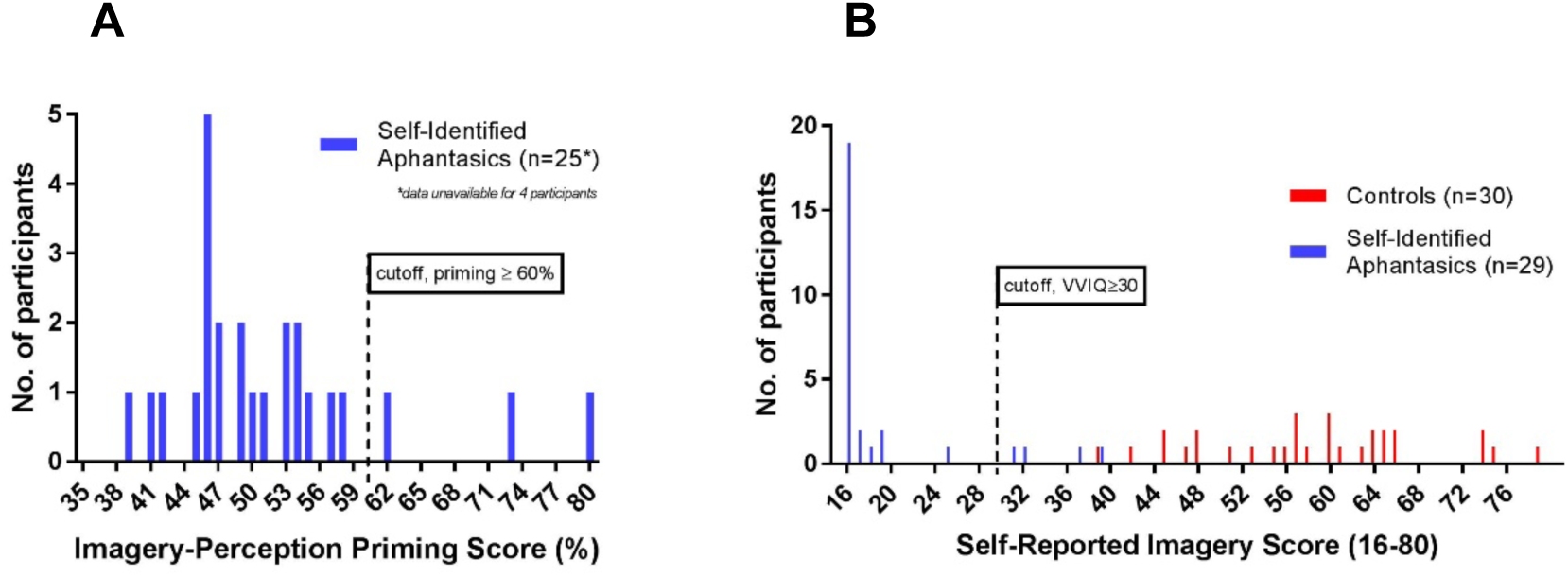
Verifying aphantasia with the VVIQ (subjective) and imagery-based priming (objective) measures of visual mental imagery. Self-identified aphantasic study volunteers’ binocular rivalry sensory strength of imagery scores. As shown, 3 self-identified aphantasic participants were excluded from the Imagery and/or Perception experiments due to imagery-based priming scores in excess of 60%, suggesting detectable sensory imagery. **B.** Self-report VVIQ scores for self-identified aphantasic study volunteers and control participants recruited for both experiments. As shown, all candidate control participants (an overlapping subset of whom was used in each experiment) received VVIQ scores in excess of 30 (out of a possible 80). The majority of self-identified aphantasic candidates reported floor-level VVIQ scores, while four were excluded from the Imagery and/or Perception experiments due to VVIQ scores in excess of 30, suggesting non-negligible subjective experience of visual mental imagery.

**Supplementary Figure S2.**
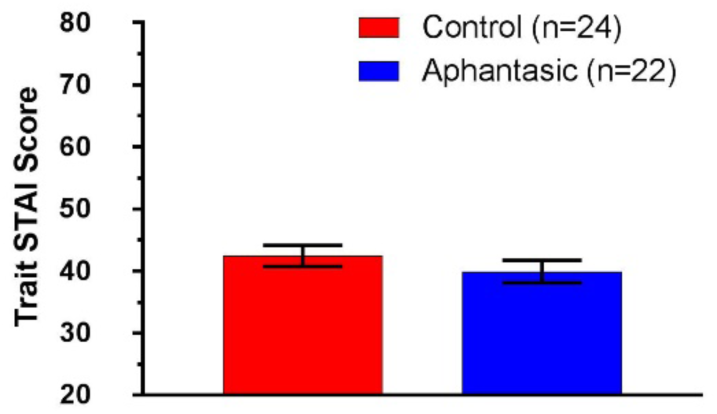
Imagery experiment, self-reported trait anxiety in aphantasic and control participants. Mean and SEM of self-reported trait anxiety measured by the STAI for aphantasic and control participants in the imagery experiment. Group averages do not significantly differ from one another (*p* = 0.32).

**Supplementary Figure S3.**
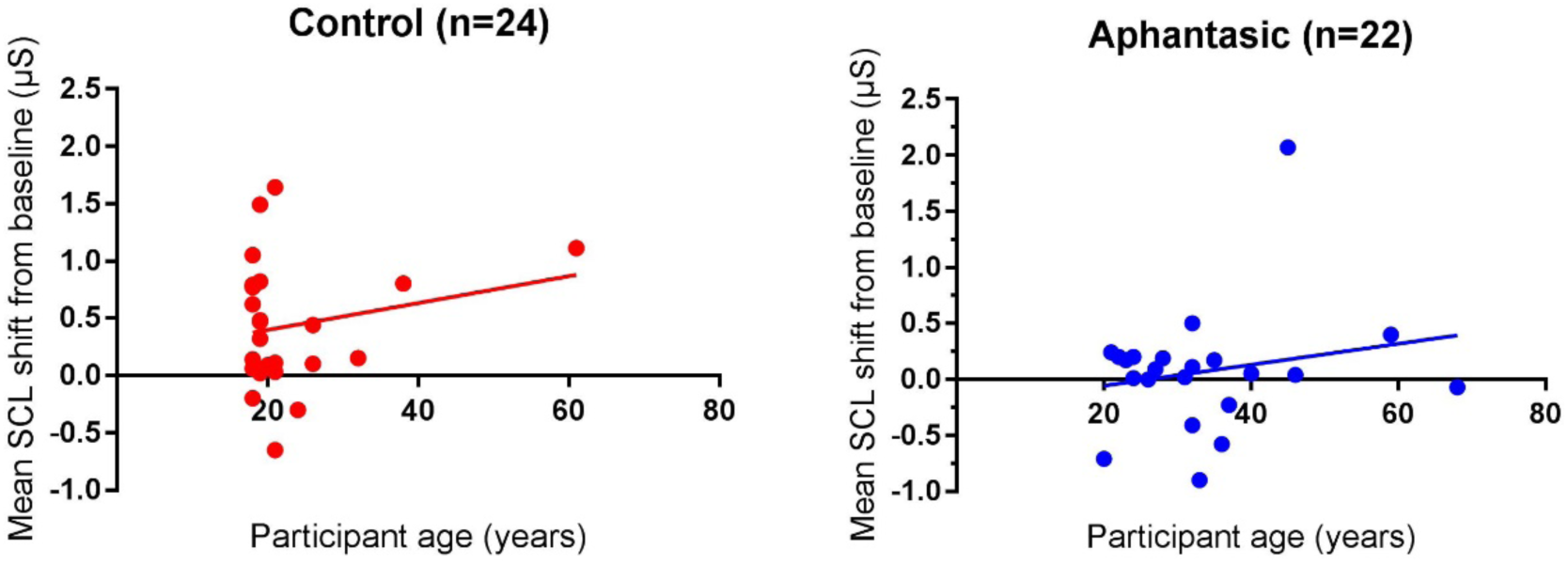
Imagery experiment, correlations between participant age and overall SCL shift from baseline while reading fictitious scenarios. Mean SCL shift from baseline for individual participants in each group in the imagery experiment, plotted against participant age. Positive regression line slopes do not significantly differ from zero in either case (aphantasic, *p*=0.37, control, *p*=0.35).

**Supplementary Figure S4.**
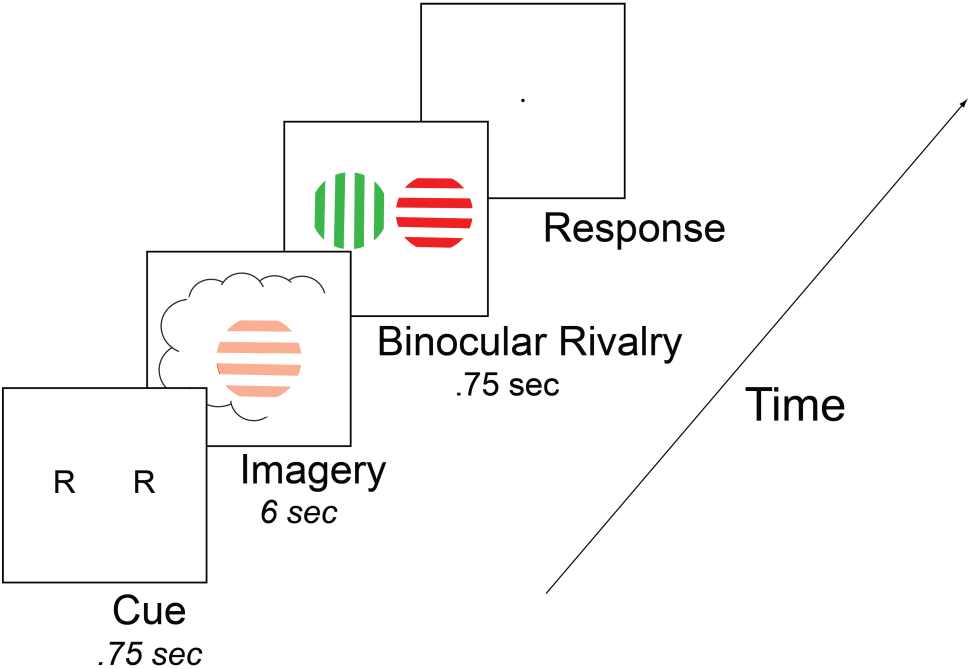
Experimental timeline for binocular rivalry imagery task.

## References

1 Schacter, D. L., Addis, D. R., & Buckner, R. L. (2007). Remembering the past to imagine the future: the prospective brain. Nature reviews neuroscience, 8(9), 657

2 Pearson J, Naselaris T, Holmes EA, Kosslyn SM. Mental Imagery: Functional Mechanisms and Clinical Applications. Trends Cogn Sci. 2015 Oct;19(10):590–602

3 Holmes, E. A., Grey, N., & Young, K. A. D. (2005). Intrusive images and “hotspots” of trauma memories in posttraumatic stress disorder: An explanatory investigation of emotions and cognitive themes. Journal of Behaviour Therapy and Experimental Psychiatry, 36, 3–17.

4 Hackmann, A., Clark, D. M., & McManus, F. (2000). Recurrent images and early memories in social phobia. Behaviour Research and Therapy, 38, 601-610; Hackman, A., Surawy, C., & Clark, D. M. (1998). Seeing yourself through others’ eye: a study of spontaneously occurring images in social phobia, Behavioural and Cognitive Psychotherapy, 26, 3-12; Hinrichsen & Clark, 2003.

5 Holmes, E. A., & Mathews, A. (2010). Mental imagery in emotional and emotional disorders. Clinical Psychology Review, 30, 349-362; Holmes, E. & Mathews, A. (2005). Mental imagery and emotion: A special relationship? Emotion, 5(4), 489–97.

6 Dils, A. T., & Boroditsky, L. (2010). Visual motion aftereffect from understanding motion language. Proceedings of the National Academy of Sciences, 107(37), 16396–16400. http://doi.org/10.1073/pnas.1009438107

7 Burbridge, D. (1994). Galton’s 100: an exploration of Francis Galton’s imagery studies. The British Journal for the History of Science, 27, 443–463.

8 Zeman, A., Dewar, M., & Della Sala, S. (2015). Lives without imagery – Congenital aphantasia. Cortex, 73, 378–80.

9 Marks, D. (1973). Visual imagery differences in the recall of pictures. British Journal of Psychology, 64, 17–24.

10 Zeman, A., Dewar, M., & Della Sala, S. (2015). Lives without imagery – Congenital aphantasia. Cortex, 73, 378–80.

11 Pearson, J., Clifford, C., & Tong, F. (2008). The functional impact of mental imagery on conscious perception. Current Biology, 18, 982–986.

12 Keogh, R. & Pearson, J. (2017). The blind mind: No sensory visual imagery in aphantasia. Cortex, in press.

13 Kreibig, S. (2010). Autonomic nervous system activity in emotion: A review. Biological Psychology, 84, 394–421.

14 e.g. Grossberg, J. M. & Wilson, H. K. (1968). Physiological changes accompanying the vizualisation of fearful and neutral situations. Journal of Personality and Social Psychology, 10, 124–133.

15 Holmes, E. A., Geddes, J. R., Colom, F., & Goodwin, G. M. (2008). Mental imagery as an emotional amplifier: application to bipolar disorder. Behaviour research and therapy, 46(12), 1251–1258.

16 Spielberger, C. D., Gorsuch, R. L., Lushene, R., Vagg, P. R., & Jacobs, G. A. (1983). Manual for the State-Trait Anxiety Inventory. Palo Alto, CA: Consulting Psychologists Press.

17 Lang, P.J., Bradley, M.M., & Cuthbert, B.N. (2008). International affective picture system (IAPS): Affective ratings of pictures and instruction manual. Technical Report A-8. University of Florida, Gainesville, FL.

18 Mathworks, Natick, 2015.

19 ADInstruments, Sydney, 2016.

20 Lang, P.J., Bradley, M.M., & Cuthbert, B.N. (2008). International affective picture system (IAPS): Affective ratings of pictures and instruction manual. Technical Report A-8. University of Florida, Gainesville, FL.

21 Brascamp, J. W., Knapen, T. H., Kanai, R., Van Ee, R., & Van Den Berg, A. V. (2007). Flash suppression and flash facilitation in binocular rivalry. Journal of vision, 7(12), 12–12.

22 Pearson, J., Clifford, C. W., & Tong, F. (2008). The functional impact of mental imagery on conscious perception. Current Biology, 18(13), 982–986.

23 Pearson, J. (2014). New directions in mental-imagery research: the binocular-rivalry technique and decoding fMRI patterns. Current Directions in Psychological Science, 23(3), 178-183.

24 Keogh R, Pearson J. The blind mind: No sensory visual imagery in aphantasia. Cortex. 2018 Aug;105:53–60

25 Pearson, J., Clifford, C. W., & Tong, F. (2008). The functional impact of mental imagery on conscious perception. Current Biology, 18(13), 982–986.

